# *In vivo* function of the lipid raft protein Flotillin 1 during CD8^+^ T cell-mediated host surveillance

**DOI:** 10.1101/403592

**Authors:** Xenia Ficht, Nora Ruef, Bettina Stolp, Federica Moalli, Nicolas Page, Doron Merkler, Ben J. Nichols, Alba Diz-Muñoz, Verena Niggli, Jens V. Stein

**Author notes:** Address correspondence: Jens V. Stein, Ph. D., Department of Oncology, Microbiology and Immunology, University of Fribourg, Ch. du Musée, 5, 1700 Fribourg, Switzerland.

## Abstract

Flotillin-1 (Flot1) is a highly conserved, ubiquitously expressed lipid raft-associated scaffolding protein. Migration of Flot1-deficient neutrophils is impaired due to a decrease in myosin II-mediated contractility. Flot1 also accumulates in the uropod of polarized T cells, suggesting an analogous role in T cell migration. Here, we analyzed morphology and migration of naïve and memory WT and Flot1^-/-^ CD8^+^ T cells in lymphoid and non-lymphoid tissues with intravital two-photon microscopy, as well as their clonal expansion during antiviral immune responses. Flot1^-/-^ CD8^+^ T cells displayed minor alterations in cell shape and motility parameters *in vivo* but showed comparable homing to lymphoid organs and infiltration into non-lymphoid tissues. Taken together, Flot1 plays a detectable but unexpectedly minor role for CD8^+^ T cell behavior under physiological conditions.

## Introduction

Leukocytes continuously recirculate through the host organism for efficient protection. As an example, naïve CD8^+^ T cells (T_N_) actively migrate in lymphoid tissue interstitium and use their unique T cell receptor (TCR) to scan dendritic cells (DCs) for cognate peptide presented on major histocompatibility complex-I (pMHC-I). Upon activation, CD8^+^ T cells undergo clonal expansion and give rise to cytotoxic effector T cells (T_EFF_). T_EFF_ spread from lymphoid organs to infected tissue where they kill host cells presenting the same cognate pMHC complex in order to eliminate the pool of intracellular pathogens, in particular viruses. After clearance of infections, long-lived memory T cells persist to provide continuous immune surveillance and rapid effector functions in the event of a secondary infection with the same pathogen. Distinct subpopulations of memory CD8^+^ T cells are categorized according to their functions and tissue homing properties: CD62L^+^ CCR7^+^ central memory T cells (T_CM_) recirculate through lymphoid organs similar to T_N_, while CD62L^-^ CCR7^-^ effector memory T cells (T_EM_) recirculate through non-lymphoid organs and blood. In recent years, a novel subset of tissue-resident memory T cells (T_RM_) were identified. T_RM_ do not recirculate but permanently reside in peripheral organs, including the epidermis and the submandibular salivary gland (SMG). In these organs, T_RM_ mediate rapid recall responses to prevent pathogen spread (1-7).

Studies using intravital twophoton microscopy (2PM) of lymphoid and non-lymphoid organs have uncovered a remarkable motility of T_N_, T_EFF_ and memory CD8^+^ T cells in all tissues analyzed thus far. This behavior is in line with their pMHC restriction, imposing the need to physically interact with DCs and target cells. Thus, the ability of CD8^+^ T cells to scan their environment through active migration is a key feature maintained throughout all phases of adaptive immune responses. Active movement of T cells requires polarization and constant cytoskeletal rearrangement – most importantly the treadmilling of fibrillar actin and the contraction of non-muscle Myosin IIa (Myo IIa) (8-12). Thus, isolated naïve T cells are round and unpolarized, but can rapidly form a characteristic polarized amoeboid shape after chemokine stimulation. This shape is characterized by a protrusive leading edge termed pseudopod and a contractile cell rear called uropod. Uropod contractility is important for detachment of uropods from adhesive substrates and for creating force to squeeze the biggest organelle of a cell, the nucleus, through narrow pores encountered during migration (8,13,14). In addition to the Myo IIa activity for actomyosin contraction, the uropod is rich in phosphorylated membrane-to-cytoskeleton-linker proteins of the ezrin/radixin/moesin family (pERM), adhesion receptors such as CD44 and PSGL-1, and cholesterol rich membrane domains, the so-called lipid rafts (15). The tip of the uropod of polarized leukocytes also contains flotillin-1 (Flot1; also known as Reggie2) and flotillin-2 (Flot2; Reggie1), evolutionary conserved, and ubiquitously expressed membrane-associated scaffolding proteins (16-21). Both flotillins possess N-terminal fatty acid modifications next to or within their prohibitin homology domain (PHB) that target them to lipid rafts (18-21). In leukocytes, C-terminal interactions lead to the hetero-oligomerization of Flot1 and −2, which is required for mutual stabilization and targeting to lipid rafts (19,22). Flotillins have been implicated in a variety of cellular functions, including cell-cell adhesion (19), endocytosis (19), regulation of G-protein coupled receptor signaling (23) and modulation of the actomyosin cytoskeleton of leukocytes. Thus, Flot1^-/-^ mice are deficient in recruitment of innate immune cells to inflammatory sites due to a decreased migratory capacity of neutrophils and monocytes (24). Flot-1^-/-^ neutrophils display reduced levels of phosphorylated myosin regulatory chain, which in turn leads to a defect in Myo IIa activity and cell migration (24). Similarly, Flot1 and Flot2 interdependently accumulate at uropods of primary human T cells, and their absence impairs uropod formation (18,22,25,26). *In vitro* experiments suggest that organization of membrane microdomains by flotillins is required for optimal T cell migration (27). Furthermore, flotillin-containing lipid rafts assemble at immunological synapses (IS) and have been proposed to serve as scaffold for the TCR signaling machinery (28-30). Yet, there is no evidence to date on how flotillins affect CD8^+^ T cell-mediated organ surveillance *in vivo*, nor how they impact T cell activation during adaptive immune responses. Here, we analyzed *in vitro* properties and *in vivo* behavior of Flot1^-/-^ CD8^+^ T cells using functional readouts under physiological conditions in mouse models. Our data suggest that Flot1 is involved in regulating the shape and speed of migrating CD8^+^ T cells in lymphoid and non-lymphoid tissues, but has only a minor impact on their ability to expand, differentiate and surveille distinct microenvironments. Taken together, our data shed light on the physiological impact of this conserved protein during adaptive immunity mediated by CD8^+^ T cells.

## Results

### In vitro characterization of naive Flot1^-/-^ T cells

Flot1 is a lipid-raft-associated protein, which accumulates at uropods of neutrophils and primary T cells (**Figure 1A**) (18). RNAseq data from the Immgen database (www.immgen.org) revealed Flot1 expression in murine CD8^+^ T cells throughout an immune response (**Figure 1B**). We decided to analyze the role of Flot1 for CD8^+^ T cell migration and activation *in vivo*. Western blot analysis confirmed the loss of Flot1 expression in Flot1^-/-^ T cells and a concomitant decrease of Flot2 as described for Flot1^-/-^ neutrophils (**Figure 1C**) (24). In the latter cell type, the remaining pool of Flot2 is cytoplasmic and not associated with lipid rafts, in line with a mutual stabilizing role for heterodimers or oligomers in leukocytes. Polyclonal Flot1^-/-^ CD8^+^ T cells isolated from lymphoid organs displayed a *bona fide* naïve CD62L^high^ CD44^low^ phenotype and showed comparable levels of surface CCR7, indicating that T_N_ were predominant in secondary lymphoid organs (SLOs) (**Figures 1D and 1E** and not shown). We compared the *in vitro* ability of naïve and activated WT and Flot1^-/-^ T cells to polarize after chemokine stimulation. Flot1 accumulated in the uropod of polarized murine T cells, comparable to findings in human T cells (**Supplemental Figure 1**). Activated Flot1^-/-^ T cell blasts but not naïve Flot1^-/-^ T cells showed a minor tendency to polarize less and to accumulate less pERM at the uropod as compared to WT blasts (**Figures 1F-H**). We next analyzed the chemotactic ability of naïve WT and Flot1^-/-^ T cells towards CCL21 across 3 µm and 5 µm pore size filters. In line with their intact polarization, both cell types migrated similarly towards CCL21 (**Figure 1I**). Since lipid raft-borne flotillin hetero-oligomers have been implicated in cortical actin binding, we hypothesized that Flot1 could influence membrane tension or membrane-to-cortex attachment. To address this issue, we measured the membrane tether force by pulling a cantilever of an atomic force microscope from the surface of activated WT and Flot1^-/-^ T cell blasts (**Figure 1J**). Lack of Flot1 did not alter the static tether force as compared to WT T cells (**Figure 1K**). In sum, lack of Flot1 has no major impact on membrane-to-cortex attachment, nor on the ability of T cells to polarize and migrate in response to homeostatic chemokines.

**Figure 1.**
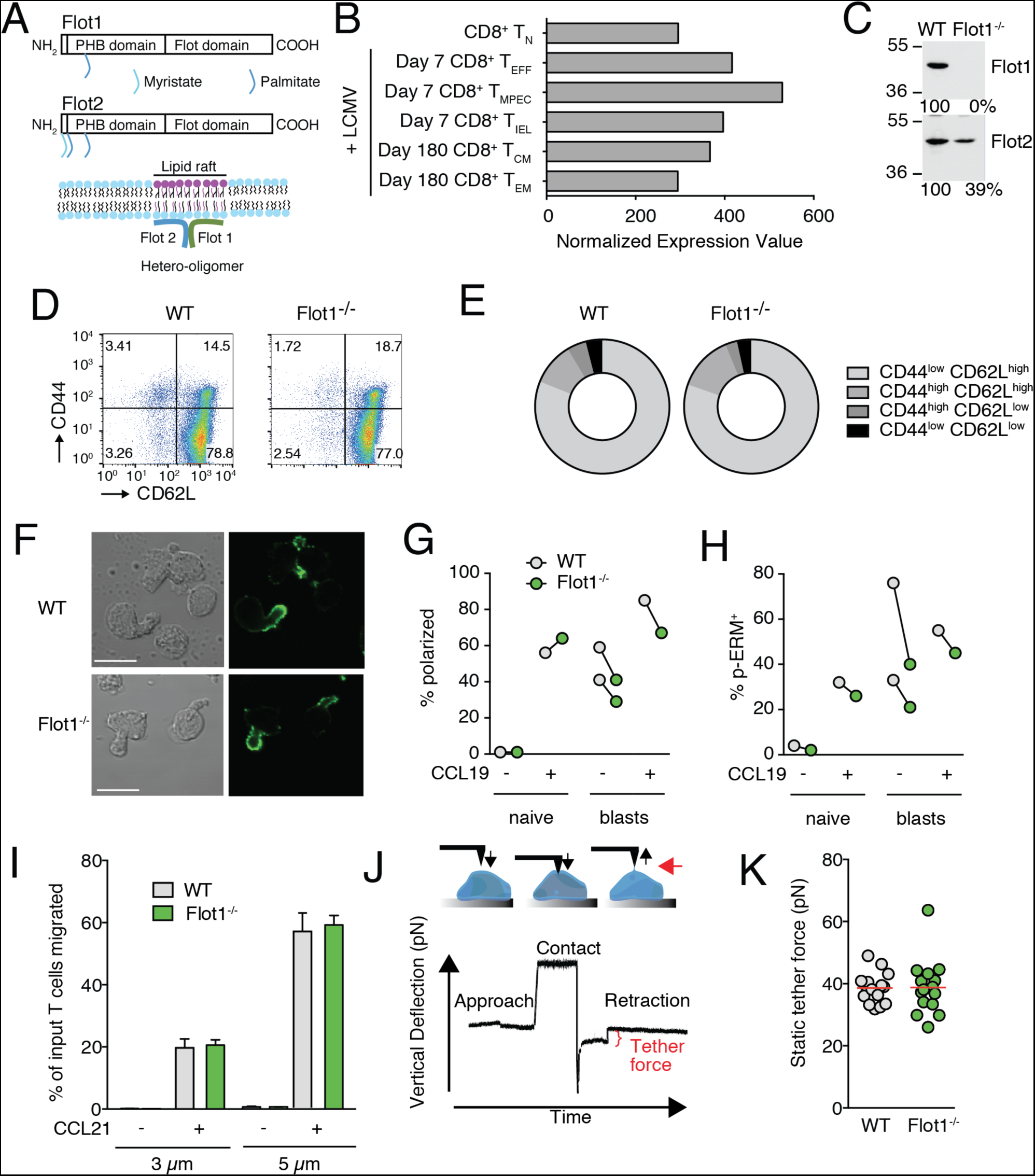
*In vitro* characterization of Flot1^-/-^ T cells. **A**. Scheme of Flot1 and 2 protein domains and association with lipid raft microdomains. PHB, prohibitin homology domain. **B.** RNA-seq data of Flot1 expression in CD8^+^ T cells (from Immgen database). **C.** Expression of Flot-1 and Flot-2 in T-lymphoblasts isolated from spleens of wild type and Flot1^-/-^ mice. **D.** Flow cytometry plot of CD62L and CD44 expression on splenic CD8^+^ T cells. **E.** Quantification of surface expression of CD44 and CD62L on CD8^+^ T cells from WT and Flot1^-/-^ mice. Pooled from 2 independent experiments, n = 4. **F.** Phase contrast and immunofluorescent images of polarized activated WT and Flot1^-/-^ T cell blasts stimulated with CCL19 and stained for pERM. Scale bar, 10 µm. **G, H.** Quantification of polarization (G) and pERM capping (H) of naïve and activated T cells with and without CCL19 stimulation. **I.** Chemotaxis of naïve WT and Flot1^-/-^ T cells towards 100 nM CCL21 through 3 and 5 µm filters. Shown is mean ± SD of one of two experiments performed in triplicates. **J.** Schematic of pulling static tethers with an atomic force microscope. **K.** Average static tether force per cell. Data shows one representative experiment of two.

### In vivo characterization of naïve Flot1^-/-^ T cell migration in lymphoid tissue

We next assessed whether Flot1 deficiency affects T cell trafficking under physiological conditions, which are difficult to reproduce in reductionist *in vitro* assays. To this end, we co-transferred WT and Flot1^-/-^ CD8^+^ T_N_ into recipient mice for homing and intravital microscopy (**Figure 2A**). We recovered comparable numbers of WT and Flot1^-/-^ CD8^+^ T cells from SLOs at 2 and 24 h after transfer (**Figure 2B** and not shown), suggesting similar homing capacity of both cell types. To analyze interstitial migration in SLOs, we co-transferred WT dsRED^+^ and Flot1^-/-^ GFP^+^ CD8^+^ T_N_ into recipient mice and performed intravital 2PM imaging of the popliteal lymph node (PLN). Consistent with the T cell homing results, we observed both cell types inside the PLN interstitium (**Figure 2C**). Time lapse imaging uncovered robust amoeboid motility WT and Flot1^-/-^ CD8^+^ T cells (**Figures 2C and 2D; Movie S1**). To quantify cell polarization, we determined the shape factor of migrating cells as described (31). A shape factor of 1 corresponds to a perfect circle, while values close to zero correlate to elongated or irregular shape outlines. Our analysis showed that Flot1^-/-^ CD8^+^ T_N_ displayed significantly rounder shapes as compared to WT CD8^+^ T cells (**Figure 2E**). This correlated with a minor but significant drop in median cell speeds (from 13.5 ± 4.0 µm/min for WT CD8^+^ T_N_ to 13.0 ± 4.1 µm/min for Flot1^-/-^ CD8^+^ T_N_; mean ± SD) (**Figure 2F**). Yet, the arrest coefficient, defined as percent of track length with speeds slower than a defined threshold value, and the meandering index were comparable between both populations (**Figures 2G and 2H**). Furthermore, both WT and Flot1^-/-^ CD8^+^ T_N_ preferentially change direction with shallow turning angles (∼ 30°) while steep turns (∼ 90°) or U-turns (∼ 180°) were rare (not shown). Both persistence and speed contribute to the motility coefficient, which measures the displacement of randomly migrating cells over time and is calculated from plotting mean displacement versus time. WT and Flot1^-/-^ CD8^+^ T_N_ display comparable motility coefficients of 54.0 and 50.9 µm^2^/min, respectively (**Figure 2I**). In sum, interstitial Flot1^-/-^ T cells are more rounded than their WT counterparts and display a minor speed drop, which did not significantly hinder the ability CD8^+^ T_N_ to scan the lymphoid microenvironment *in vivo*.

**Figure 2.**
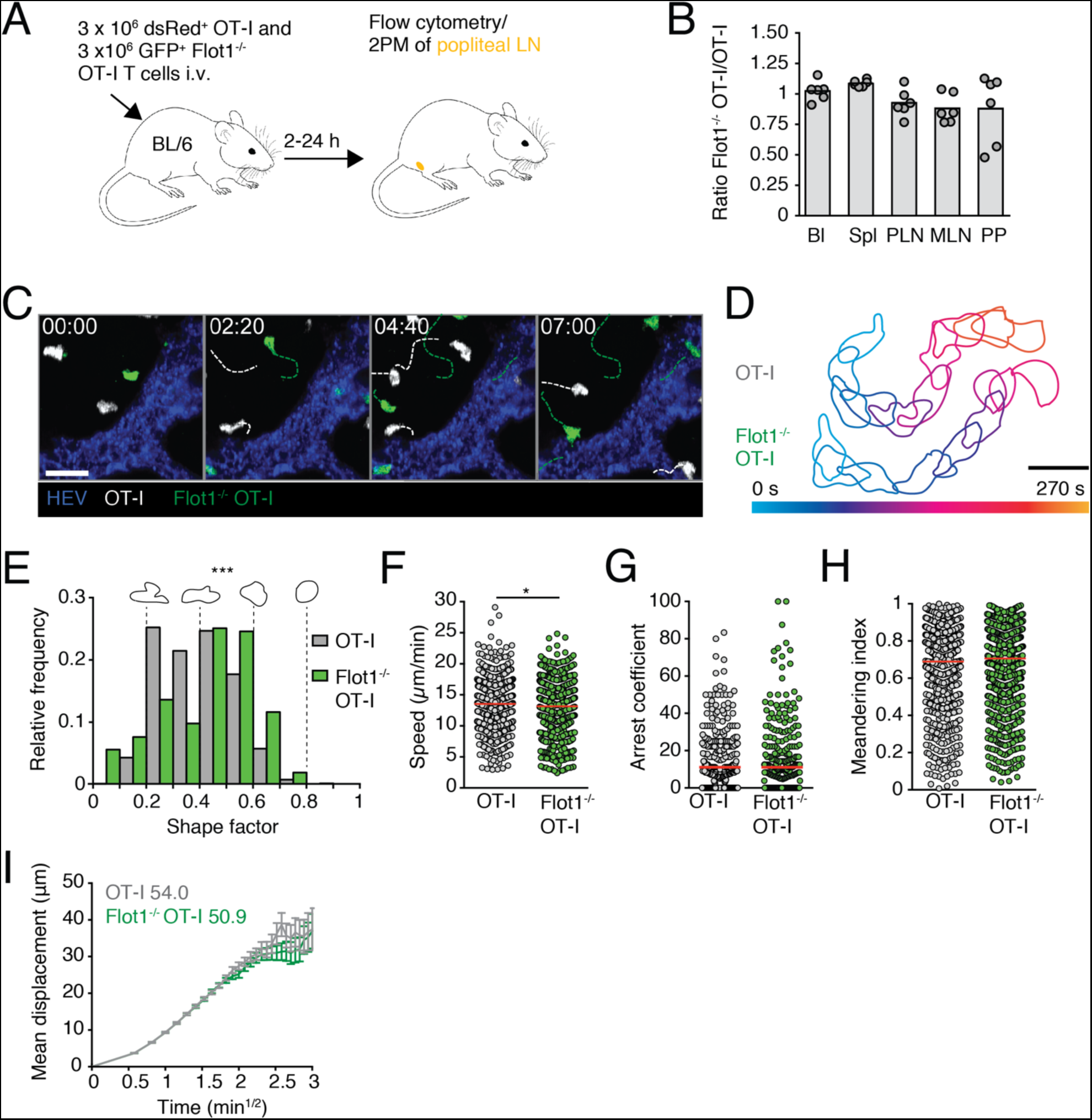
*In vivo* trafficking of naïve Flot1^-/-^ CD8^+^ T cells in lymphoid organs. **A.** Experimental layout. **B.** 2 h *in vivo* homing of OT-I WT or Flot1^-/-^ OT-I T cells to indicated organs: blood (Bl), spleen (Spl), peripheral lymph nodes (PLN), mesenteric lymph nodes (MLN) and Peyer’s patches (PP). Each dot represents organs from one mouse, bars depict mean. Pooled from 2 independent experiments with a total of 6 mice per group. **C.** 2PM image sequence of naive WT and Flot1^-/-^ OT-I T cells migrating in popliteal LN. Dashed white and green lines indicate tracks of migrating WT and Flot1^-/-^ OT-I T cells, respectively. Scale bar, 20 µm; Time in min:s. **D.** Representative time-coded outlines of migrating WT and Flot1^-/-^ OT-I T cells. Scale bar, 10 µm. **E.** Shape factor of naive WT and Flot1^-/-^ OT-I T cells migrating in popliteal LN. Pooled from 2 independent experiments with 3 mice per group. **F-H.** Speed (F), arrest coefficient (G) and meandering index (H) of naïve WT or Flot1^-/-^ OT-I T cells migrating in popliteal LN. Red lines depict median, each dot represents the average value for an individual track. **I.** Mean displacement versus time for datasets in F-H. Numbers indicate motility coefficient in μm^2^/min. Data in F-I are pooled from 3 independent experiments with a total of 8 mice. Data in B were tested for significance with 2-way ANOVA and Sidak’s multiple comparison test. Data in F were analyzed using unpaired Student’s t-test, data in E, G and H using Mann-Whitney test. *, p < 0.05; ***, p < 0.001.

### Activation, expansion and memory formation of Flot1^-/-^ CD8^+^ T cells

Flotillin-rich lipid rafts have been implicated in TCR signaling at the IS, contributing to T cell activation *in vitro* (28-30). In contrast, activation and memory formation of Flot1^-/-^ CD8^+^ T cells has not been systematically analyzed *in vivo* to date. To explore this further, we adoptively transferred either GFP^+^ WT or GFP^+^ Flot1^-/-^ OT-I T cells into recipient mice. OT-I T cells recognize the amino acid sequence SIINFEKL derived from Ovalbumine (OVA) (32). At 24 h post transfer, we induced a systemic infection by i.p. injection of OVA-encoding Lymphocytic Choriomeningitis Virus (LCMV-OVA) (**Figure 3A**) (33). We observed a comparable clonal expansion of WT and Flot1^-/-^ OT-I T cells during the acute phase (day 6 p.i.), followed by a contraction after the effector phase (day 15 p.i.) and the formation of a long-lived memory pool (> day 28 p.i.) in lymphoid tissue and SMG (**Figure 3B**). In the effector phase, activated CD8^+^ T cells differentiate from KLRG-1^-^ CD127^-^ early effector cells (EEC) into KLRG-1^+^ CD127^-^ short-lived effector cells (SLEC) or KLRG-1^-^ CD127^+^ memory precursor cells (MPEC), with few KLRG-1^+^ CD127^+^ double-positive effector cells (DPEC) (34). The decision of becoming SLEC versus MEPC is influenced by the strength of antigenic stimulation and the cytokine milieu and serves therefore as readout for alterations in CD8^+^ T cell responsiveness. When we examined WT and Flot1^-/-^ OT-I T cells on day 6 p.i., we observed most cells to be KLRG-1^-^ CD127^-^ EECs or KLRG-1^+^ CD127^-^ SLECs without notable changes in population ratios between WT or Flot1^-/-^ OT-I T cells (**Figures 3C and 3D**), indicating comparable activation.

**Figure 3.**
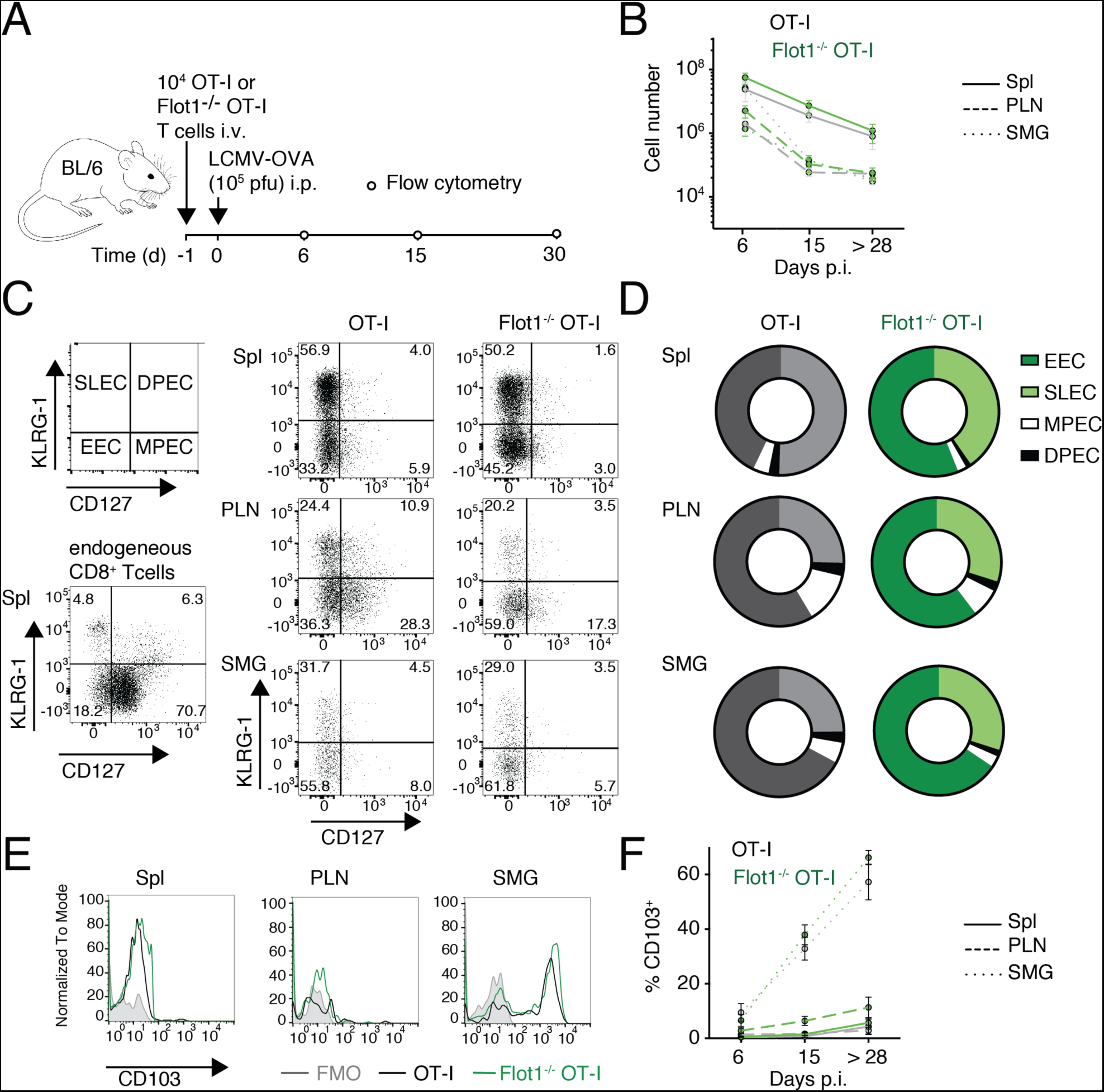
Expansion and memory formation of Flot1^-/-^ CD8^+^ T cells. **A.** Experimental layout of LCMV-OVA infection. **B.** Total number of GFP^+^ cell number in spleen (Spl), pooled peripheral lymph nodes (PLN) and submandibular salivary gland (SMG) at day 6, 15, and >28 p.i. Pooled from 2 independent experiments with a total of 5-6 mice per group. **C, D.** Representative flow cytometry plots (C) and pooled results (D) of CD127 and KLRG-1 co-staining of endogenous CD8^+^ and transferred OT-I T cells on 6 d p.i.. Pooled from 2 independent experiments and a total of 4-8 mice. **E, F.** Representative flow cytometry plots (E) and pooled results (F) of CD103 expression on WT and Flto1^-/-^ OT-I T cells in memory phase (>28 days p.i.). F shows mean with SEM from 2-4 independent experiments with a total of 5-12 mice per group and time point.

Finally, we analyzed whether T_RM_ formation in SMG is affected by absence of Flot1. Both WT or Flot1^-/-^ OT-I T cells exhibited a comparable gradual increase of the T_RM_ marker CD103 in SMG but not PLN or spleen, suggesting largely preserved capacity to form *bona fide* T_RM_ in non-lymphoid organs (**Figures 3E and 3F**). Taken together, our data suggest normal clonal expansion, differentiation, tissue infiltration and memory formation of Flot1^-/-^ CD8^+^ T cells during an antiviral immune response.

### In vivo migration of Flot1^-/-^ CD8^+^ memory T cells in lymphoid tissue

We set out to examine the tissue surveillance properties of memory CD8^+^ T cells in lymphoid tissue after viral infections. To this end, we co-transferred dsRed^+^ WT and GFP^+^ Flot1^-/-^ OT-I CD8^+^ T cells into recipient mice, which were infected with LCMV-OVA one day later. After > day 30 p.i., we performed intravital imaging of PLN (**Figure 4A**). Both WT and Flot1^-/-^ OT-I T cells displayed active amoeboid migration in the PLN interstitium (**Figures 4B and 4C; Movie S2**), similar to T_N_. Compared to WT memory OT-I T cells, Flot1^-/-^ T cells were on average rounder than their WT counterparts (**Figure 4D**). This correlated to a minor, but statistically significant reduction in migration speeds of memory Flot1^-/-^ OT-I T cells as compared to WT OT-I T cells (from 11.2 ± 3.9 µm/min for WT memory CD8^+^ T cells to 10.2 ± 3.7 µm/min for Flot1^-/-^ memory CD8^+^ T cells; mean ± SD) (**Figure 4E**). In addition, the arrest coefficient was increased in Flot1^-/-^ memory OT-I T cells, while the meandering index of Flot1^-/-^ memory T cells was slightly reduced as compared to WT OT-I T cells (**Figures 4F and 4G**). As a consequence, Flot1^-/-^ memory T cells displayed a reduced motility coefficient compared to WT memory T cells (**Figure 4H**). In sum, Flot1 plays a detectable role in the morphology and migration of lymphoid tissue memory CD8^+^ T cells, which mostly consist of T_CM_. Nonetheless, the ability of CD8^+^ T cells lacking Flot1 to scan the lymphoid microenvironment remains comparably robust.

**Figure 4.**
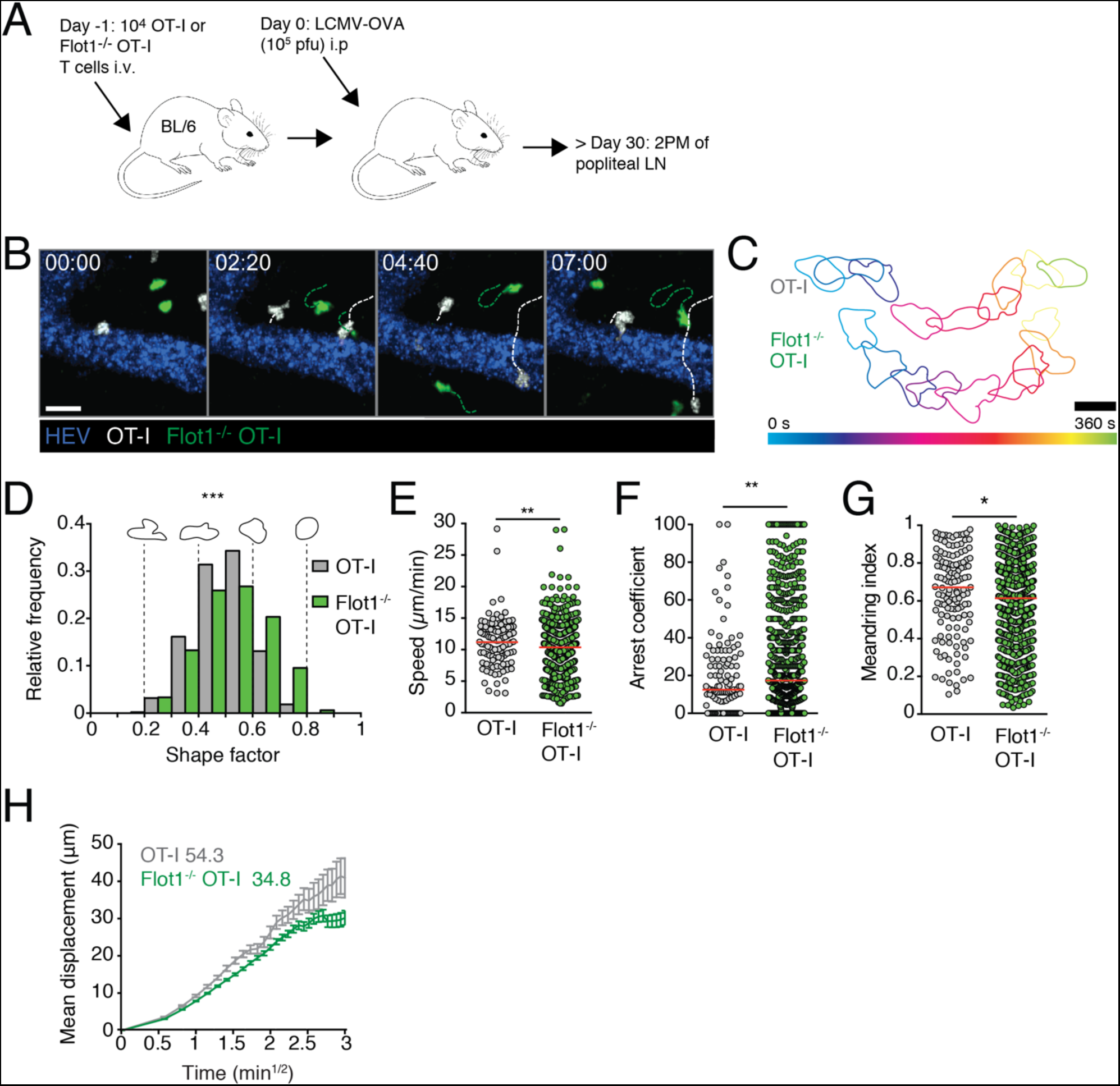
*In vivo* trafficking of memory Flot1^-/-^ CD8^+^ T cells in lymphoid organs. **A.** Experimental layout. **B.** 2PM image sequence of memory WT and Flot1^-/-^ OT-I T cells migrating in popliteal LN. Dashed white and green lines indicate tracks of migrating memory WT and Flot1^-/-^ OT-I T cells, respectively. Scale bar 20 µm; Time in min:s. **C.** Representative time-coded outlines of migrating memory WT and Flot1^-/-^ OT-I T cells in PLN. Scale bar, 10 µm. **D.** Shape factor of memory WT and Flot1^-/-^ CD8^+^ T cells migrating in popliteal LN. Pooled from 2 independent experiments with 3 mice per group. **E-G.** Speed (E), arrest coefficient (F) and meandering index (G) of memory WT or Flot1^-/-^ OT-I T cells migrating in LN. Red lines depict median, each dot represents the average value for an individual track. **H.** Mean displacement versus time for datasets in E-G. Numbers indicate motility coefficient in μm^2^/min. Flot1^-/-^ OT-I data in E-I are pooled from 3 independent experiments (n = 8) and WT OT-I data are pooled from 2 independent experiments (n = 5). Data in E were analyzed using unpaired Student’s t-test, data in D, F and G using Mann-Whitney test. *, p < 0.05; **, p < 0.01; ***, p < 0.001.

### Patrolling behavior of Flot1^-/-^ CD8^+^ T_RM_ in epidermis

Since uropod function is critical in dense environments for nuclear propulsion and Flot1 is required for neutrophil accumulation in non-lymphoid tissue (11,24,35,36), we reasoned that the function of Flot1-mediated control of cell shape may become more apparent in dense non-lymphoid tissue. To address this point experimentally, we co-transferred GFP^+^ Flot1^-/-^ OT-I and dsRed^+^ WT OT-I T cells into WT recipients and performed skin tattooing of a OVA-encoding Herpes Simplex Virus I (HSV-OVA) one day later as described (**Figure 5A**) (6). In this setting, viral infections lead to the recruitment of CD8^+^ effector T cells, which differentiate after viral clearance into epidermal T_RM_ (33). At 5 weeks p.i., we surgically prepared the tattooed skin for intravital imaging of WT and Flot1^-/-^ T_RM_. Epidermal T_RM_ are located within the basal epidermis above a loose layer of dermal collagen, which can be visualized in 2PM by second harmonic generation. Owing to the dense epithelial microenvironment, epidermal T_RM_ migrate at 1-2 μm/min while displaying a highly dendritic morphology with many finger-like protrusions (37,38). We observed that both WT and Flot1^-/-^ epidermal T_RM_ exhibit this typical morphology and localization (**Figures 5B and 5C; Movie S3**). Shape factors of both WT and Flot1^-/-^ epidermal T_RM_ were much lower than of lymphoid tissue T cells, reflecting their highly elongated morphology. In contrast to Flot1^-/-^ T cells in lymphoid organs, epidermal Flot1^-/-^ T_RM_ displayed a significantly more elongated morphology as compared to WT cells (**Figures 5C and 5D**). WT epidermal T_RM_ migrated with a mean speed of 1.7 ± 0.7 μm/min, while Flot1^-/-^ epidermal T_RM_ migrated with a reduced speed of 1.4 ± 0.6 μm/min (mean ± SD) (**Figure 5E**). Furthermore, Flot1^-/-^ T_RM_ had a higher arrest coefficient, but a similarly low meandering index as WT T_RM_ (**Figures 5F and 5G**). Despite these minor changes in cell motility parameters, the motility coefficient of epidermal Flot1^-/-^ epidermal T_RM_ was unchanged compared to WT T_RM_ (**Figure 5H**). In conclusion, Flot1^-/-^ epidermal T_RM_ retain the capacity of scanning even a very dense microenvironment consisting of epithelial layers.

**Figure 5.**
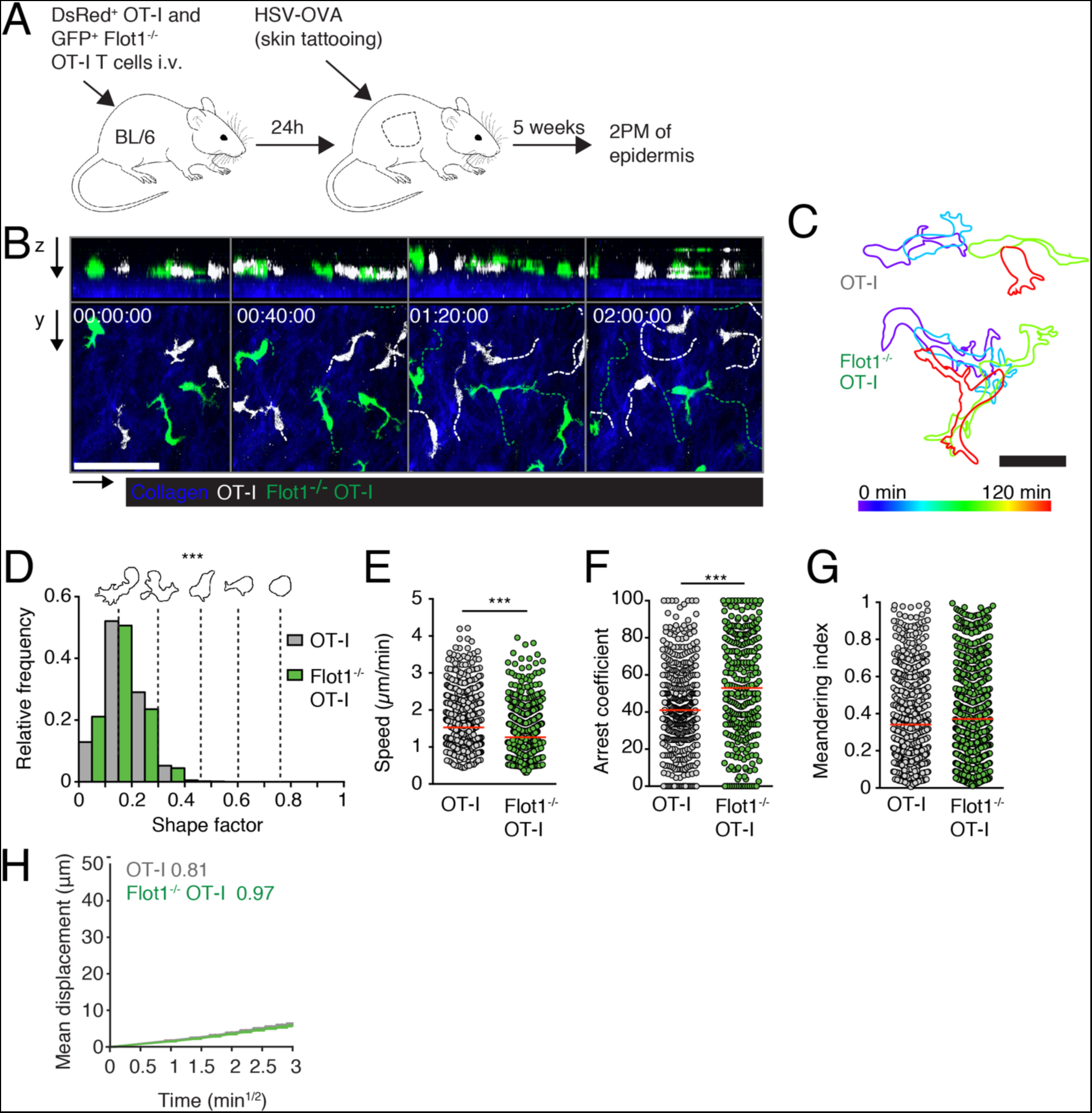
Epidermal Flot1^-/-^ CD8^+^ T_RM_ migration. **A**. Experimental layout for the generation of epidermal T_RM_ with HSV-OVA infection. **B.** 2PM image sequence of memory WT and Flot1^-/-^ OT-I T cells showing both xy and xz views of epidermis over 2 h of continuous imaging. WT OT-I cells and cell tracks in white, and Flot1^-/-^ OT-I cells and tracks in green. Dermal collagen visualized by second harmonic generation shown in blue. Scale bar 50 µm; Time in h:min:s. **C.** Representative time-coded outlines of migrating memory WT and Flot1^-/-^ OT-I T cells in the epidermis. Scale bar, 20 µm. **D.** Shape factor of epidermal WT and Flot1^-/-^ OT-I T cells. Pooled from one experiment with n = 2 (WT OT-I) or n = 3 (Flot1^-/-^ OT-I) mice. **E-G.** Speed (E), arrest coefficient (F) and meandering index (G) of epidermal WT or Flot1^-/-^ OT-I T cells. Red lines depict median, each dot represents the average value for an individual track. **H.** Mean displacement versus time for datasets in E-G. Numbers indicate motility coefficient in μm^2^/min. Data in E to H were pooled from 3 independent experiments (n = 10). Data in E were analyzed using unpaired Student’s t-test, data in D, F and G using Mann-Whitney test. ***, p < 0.001.

## Discussion

CD8^+^ T cells are remarkably efficient in physically scanning a wide variety of target tissues in search of pMHC-I-presenting DCs and target cells. To this end, these cells acquire a polarized shape with a protrusive leading edge and a contractile uropod. Flotillins are highly conserved proteins that scaffold lipid rafts and control the cortical cytoskeleton of neutrophil uropods, where Flot-rich membrane domains accumulate (24). Yet, their *in vivo* function for adaptive immune cells has not been explored to date. Our data suggest a minor role for Flot1 in optimizing CD8^+^ T cell speeds in lymphoid and non-lymphoid tissues. Furthermore, Flot1 contributes to shape maintenance but does not affect expansion during an immune response under the experimental conditions used here.

Flotillins are involved in TCR recycling and signaling at the IS (28,30). Yet, in our system using a transgenic TCR (OT-I) and cognate peptide expressed by replication-competent viruses, we did not observe a defect in expansion and memory formation of Flot1^-/-^ CD8^+^ T cells. We assessed T cell numbers and phenotypes at day 6 post infection, which may have been too late to detect minor alterations during early T cell activation. It is also possible that the strong affinity of OT-I to its cognate peptide-MHC is sufficient to override any Flot1-mediated defects or that T cells compensate for lower signaling strength by a longer interaction time with DCs.

We used 2PM imaging of PLN and skin to asses morphology and migratory patterns of Flot1^-/-^ CD8^+^ T cells in *vivo*. We consistently found changes in cell shape: Lymphoid tissue Flot1^-/-^ T _cells_ were slightly rounder than their WT counterparts, while the opposite was observed for epidermal T_RM_. These changes are consistent with a minor decrease in contractility in the absence of Flot1. Lymphoid tissue T cells exhibit amoeboid migration, for which they require contractility to establish a polarized cell shape including the pinched middle and uropod. Flot1 therefore contributes to uropod formation in lymphoid tissue. In contrast, epidermal T_RM_ use a finger-like migration mode. It was hypothesized that actin-polymerization is driving the extension of the finger, while myosin mediated contractility is required for contraction of the protrusion and for cell translocation (39). Therefore, a minor reduction in contractility would correlate to hyperelongated protrusions in epidermis.

RNAseq expression levels of Flot1 and lipid raft content are comparable between naïve and memory CD8^+^ T cells (40). However, multiple interacting partners of flotillins, such as the adhesion molecule CD44, are differentially expressed in naïve versus memory CD8^+^ T cells. Changes in the interactome of Flot1 may therefore underlie the larger impact of Flot1 deficiency on lymph node memory CD8^+^ T cell motility as compared to T_N_. Similarly, Flot1^-/-^ epidermal T_RM_ migrated with a slower velocity compared to WT T_RM_. Presumably a small reduction in myosin-mediated contractility cannot be fully compensated in the extremely dense environment of the epidermis. Despite these minor changes in cell shape and motility parameters, Flot1 appears to play a much less prominent role in CD8^+^ T cells as compared to neutrophils (24). This may be due to the fact that while readily detectable by gene expression and western blot, Flot1 expression is several times lower in T cells as compared to neutrophils (www.immgen.org).

In sum, the present data demonstrate that Flot1^-/-^ CD8^+^ T cells are capable of expansion and formation of effector and memory T cells following viral infection. Furthermore, our imaging experiments show that morphology and homeostatic immune surveillance by Flot1^-/-^ CD8^+^ T_N_ and memory T cells in lymphoid and non-lymphoid tissue is slightly affected but not critically impaired. Thus, while flotillins are highly conserved and universally expressed, we demonstrate here that they play only minor roles for CD8^+^ T cell-mediated immune surveillance.

## Experimental Procedures

### Mice

OT-I TCR transgenic mice (32) crossed to Tg(UBC-GFP)30Scha “Ubi-GFP” (41) (“GFP^+^ OT-I”) or hCD2-dsRed (42) (“DsRed^+^ OT-I”) mice on the C57BL/6J background have been described (33). GFP^+^ OT-I mice were crossed with Flot1^-/-^ mice (24). All mice were bred at our animal facility (Bern, Switzerland) and used as lymphocyte donor mice at 6-20 weeks of age. 6-10 week-old sex-matched C57BL/6J mice (Janvier, France) were used as recipient mice. All experiments were performed in accordance to federal animal experimentation regulations and approved by the cantonal committee.

### Reagents

Sodium Pyruvate 100mM (#11360-039), HEPES buffer 1M (#15630-056), RPMI 1640 (#21875-034) L-Glutamine 200nM (#25030-024), Minimum essential Medium Non-essential amino acids (MEM NEAA, #11140-035), Pen Strep (#15140-122) and were purchased from Gibco and Fetal Bovine Serum (FCS, #SV30143.03) was obtained from HyClone.

### Cell polarization and pERM capping

Cell polarization and pERM capping experiments were performed as described (26). In brief, naïve T cells were isolated from spleen and purified by negative selection. Cells were then incubated for 15 min without or with 200 ng CCL19/ min at 37°C in suspension followed by fixation with 10% trichloroacetic acid (TCA), staining for pERM and evaluation of cell morphology and pERM caps. Only cells with pERM staining were evaluated (100 cells counted per sample). For activated T cells, T cells were isolated from spleen and incubated for 2 days with 1 µg/ml ConA, then 4 days with Il-2 (100 Units/ml). Cells were then incubated for 15 min without or with 200 ng CCL-19/ min at 37°C in suspension followed by fixation with 10% TCA, staining for pERM and evaluation of cell morphology and pERM caps. Only cells with pERM staining were evaluated (100 cells counted per sample).

### Western Blots

Expression of flotillin-1 and -2 was measured in T lymphoblasts isolated from spleens of WT and Flot1^-/-^ mice. To this end, 1.5 x 10^6^ T lymphoblasts per sample were precipitated with TCA and separated on 10% sodium dodecyl sulfate polyacrylamide gel electrophoresis (SDS-PAGE) followed by immunoblotting with the specified antibodies (anti-mouse flotillin 1 #610820 and anti-mouse flotillin 2 #610383 from BD Transduction Laboratories).

### T cell transfer and viral infections

Negative isolation of CD8^+^ T cells from peripheral lymph nodes and spleen of dsRed^+^ or GFP^+^ OT-I or Flot1^-/-^ GFP^+^ OT-I mice was performed using EasySep^TM^ Mouse CD8^+^ T cell Isolation Kit (Stem Cell Technologies). We confirmed CD8^+^ T cell purity to be >90% by flow cytometry and transferred 10^4^ or 0.5-1 x 10^6^ OT-I cells in CM-R (RPMI/10% FCS/1% HEPES/1% PenStrep/2 mM L-Glutamine/1 mM Na-Py) i.v. 1 day before LCMV-OVA or HSV-OVA infection, respectively. For some experiments co-transfer of equal amounts of Flot1^+/+^ OT-I GFP or Flot1^-/-^ OT-I GFP and Flot1^+/+^ OT-I dsRed was performed. Mice were either infected with 1 x 10^5^ pfu. LCMV-OVA (43) i.p. or anesthetized, shaved, and tattooed with 5 x 10^5^ pfu in 10 μL of HSV_TOM-OVA_ using a Cheyenne tattooing machine as described (6). Experimental read-outs for the acute, cleared and memory phase of LCMV infection were performed 6 days, 15 days and > than 28 days post infection, respectively. Experimental read-outs for memory phase of HSV infection were performed later than 35 days post infection.

### Flow cytometry

At indicated time points, spleens and PLN were harvested and and organs were passed through cell strainers (70 µm; Bioswisstec) to obtain single cell suspensions. Red blood cell lysis was performed on splenocytes. SMG was minced and treated with 2 U/µl collagenase II (Worthington Biochem) in CM-R for 45 min at 37°C, passed through a 70 µm cell strainer and washed with PBS/10 mM EDTA. We used the following reagents for flow cytometry:

| Antibody | clone | company | Order number |
| --- | --- | --- | --- |
| anti- CD4-PE | GK1.5 | Biolegend | 100408 |
| anti-CD8a-PerCP | 53-6.7 | Biolegend | 100732 |
| Anit-CD8a-APC | 53-6.7 | Biolegend | 100712 |
| anti- CD103-Biotin | M290 | Biolegend | 103106 |
| anti-CD103-APC | 2E7 | Biolegend | 121413 |
| anti- CD45-PE | 30-F11 | BD Biosciences | 550261 |
| anti-CD44-PE | IM7 | BD Biosciences | 553134 |
| Streptavidin-BV421 | - | Biolegend | 4055225 |
| Streptavidin-PE | - | Biolegend | 405203 |
| anti-CD62L-APC | MEL-14 | Biolegend | 104412 |
| anti-KLRG1-PECy7 | 2F1/KLRG1 | Biolegend | 138415 |
| anti-KLRG1-PE | 2F1/KLRG1 | Biolegend | 138407 |
| anti-CD127-Biotin | B12-1 | BD Pharmingen | 555288 |
| anti-CD127-APC | A7R34 | Biolegend | 135012 |

Cell suspensions were stained on ice for 20 min with indicated antibodies and washed in FACS buffer (PBS/2% FCS/1 mM EDTA). For acute phase stainings, cells were fixed for 20 min in 1% PFA after staining. Cells were washed again prior to acquisition and at least 10^5^ cells in the FSC/SSC “lymphocyte” gate were acquired using a FACSCalibur or LSR II (BD Bioscience). Total cell counts were measured by acquiring single cell suspensions in PKH26 reference microbeads (Sigma), or Beckman coulter counting beads (REF7547053) for 1 min at high speed or by counting with FastRead counting slides (Immune Systems Ltd). Gating for CD103, CD127 and KLRG1 was set according to isotype or FMO controls. Data was analyzed with FlowJo (FlowJo LLC).

### T cell homing

DsRed^+^ OT-I and GFP^+^ Flot1^-/-^ CD8^+^ T cells mice were injected i.v. into sex-matched recipients (2-3 x 10^6^ cells/mouse). After 2 h, mice were killed and perfused with PBS. Peripheral LN, mesenteric LN, Peyer’s Patches, and spleen were harvested and passed through passed through a 70 µm cell strainer. Afterwards, erythrocytes in spleen were lysed in RBC buffer for 5 minutes. Blood was harvested by taking it up directly from the vena cava during perfusion into a 20 ml syringed filled with 10 mL of PBS/10 mM EDTA. Samples were stained on ice with PerCP anti-mouse CD8a, washed and acquired on a FACSCalibur flow cytometer (BD). Ratio of KO to WT cells in each organ was normalized to input.

### Transwell migration assays

OT-I WT and Flot1^-/-^ CD8^+^ T cells (5 x 10^5^/well) in 100 μL CM-R were added to the inserts of 3 or 5 μm-pore size Transwell Chambers (Costar) and allowed to migrate towards the indicated concentrations of recombinant CCL21 (Preprotec) for 2 h at 37°C. Percent of transmigrated cells was measured by comparing cell counts of input to transwells after 1 min of acquisition at the same speed with a FACSCalibur flow cytometer (BD).

### Multiphoton intravital imaging

Multiphoton intravital imaging of the popliteal LN and skin was performed as previously described (33). In short, ketamine/xylazine/acepromazine was used to anesthetize mice and the right popliteal lymph node was surgically prepared. For skin imaging, hair on the right flank was removed and a section of skin was elevated onto a metal holder by making two parallel incisions. Prior to imaging, Alexa 633-conjugated MECA-79 (10 µg/mouse) was injected i.v. to mark HEV in LN or 400 – 600 µg of 70 kDa TexasRed Dextran to label all blood vessels. 2PM imaging was carried out with a TrimScope I 2PM microscope using a 25X Nikon (NA 1.0) or 20X Olympus (NA 0.95) objective (LaVisionBiotec). ImSpector software was used to control the 2PM system and acquire images. Some image series were obtained with the help of an automated system providing real-time drift correction (44). Excitation was provided by a Ti:sapphire laser (Mai Tai HP, Spectraphysics) tuned to 780 or 840 nm. For analysis of cell migration, 11 to 20 x-y slices with a z-step size of 2-4 µm (22-64 µm depth) were acquired. Frame rate was 3 per min for LN or 1 per min for skin. Second harmonic signals and emitted light were detected with 447/55-nm, 525/50-nm, 593/40-nm and 655/40-nm bandpass filters using non-descanned detectors. Image series were rendered into four-dimensional videos with Imaris (Bitplane) or Volocity (PerkinElmer) software. Tissue movement was corrected by either the correct drift function of Imaris or with a MATLAB script identifying the 3D movement in a reference channel. Motility parameters such as angle changes were derived from x, y, and z coordinates of cell centroids using Imaris, Volocity and MATLAB protocols. The cutoff for the arrest coefficient in LN was 5 µm/min and in epidermis 1 µm/min. Motility coefficients, which are a measure of a randomly moving cell’s ability to move away from its starting position, were calculated from the slope of a graph of the average mean displacement against the square root of time. Shape factor was calculated by analyzing 2D projections of cell tracks of horizontally moving cells in Volocity as described (31). For some depicted image series, raw cell signals were masked with Imaris to hide autofluorescence. Brightness and contrast were adjusted in all images.

### Membrane tension measurements

CD8^+^ T cells were isolated with Stem Cell negative selection kit as described above, and kept on ice in CM-R with 5 μg/mL IL-7 (407-ML-005 R&D) until o.n. activation with 2.5 μg/mL ConA. Imaging dishes were washed sonicated for 5 min in water, washed in 99% ethanol, dried, and sterilized with UV-light for 15 min. Dishes were coated for 30-60 min at 37°C with 0.01% Poly-L-Lysine (Sigma Aldrich), and washed 8-10 times with PBS. Approximately 5 x 10^5^ T cells were plated onto the coated dish in 0.5 mL CM-R and incubated for 10-15 min at 37°C. Subsequently, dishes were washed carefully 8-10 times with warm medium. For tether measurements, 3 mL CM-R (5% FCS) were added to the dish. A cantilever was coated with 4 mg/mL ConA for 60 min, washed and its spring constant calibrated in air. Measurements were performed with a CellHesion 200 from JPK. Analysis was performed with JPK Data Processing Software.

### Statistical analysis

As indicated in the figure legends, unpaired Student’s test, Mann-Whitney test, or Two-way ANOVA with a Sidak’s multiple comparison test were used to determine statistical significance (Prism 7, GraphPad). Significance was set at p < 0.05.

## Ethics Statement

All animal experiments were approved by the Cantonal Committee for Animal Experimentation and carried out in accordance with federal guidelines.

## Author Contributions

XF performed imaging, AFM, transwell, and flow cytometry experiments. NR performed AFM experiments. BS performed chemotaxis and flow cytometry experiments. FM assisted with skin imaging. VN performed western blots and measured pERM capping. NP and DM provided vital material. ADM provided critical assistance for membrane tension measurements. BJN generated the Flot1^-/-^ mouse line. XF and JSV designed experiments and wrote the manuscript.

## Acknowledgements

This work was funded by Swiss National Foundation (SNF) project grants 31003A_135649, 31003A_153457 and 31003A_172994 (to JVS), and Leopoldina fellowship LPDS 2011-16 (to BS). This work benefitted from optical setups of the Microscopy Imaging Center of the University of Bern.

## Supplemental Figure Legend

**Supplementary Figure 1.**
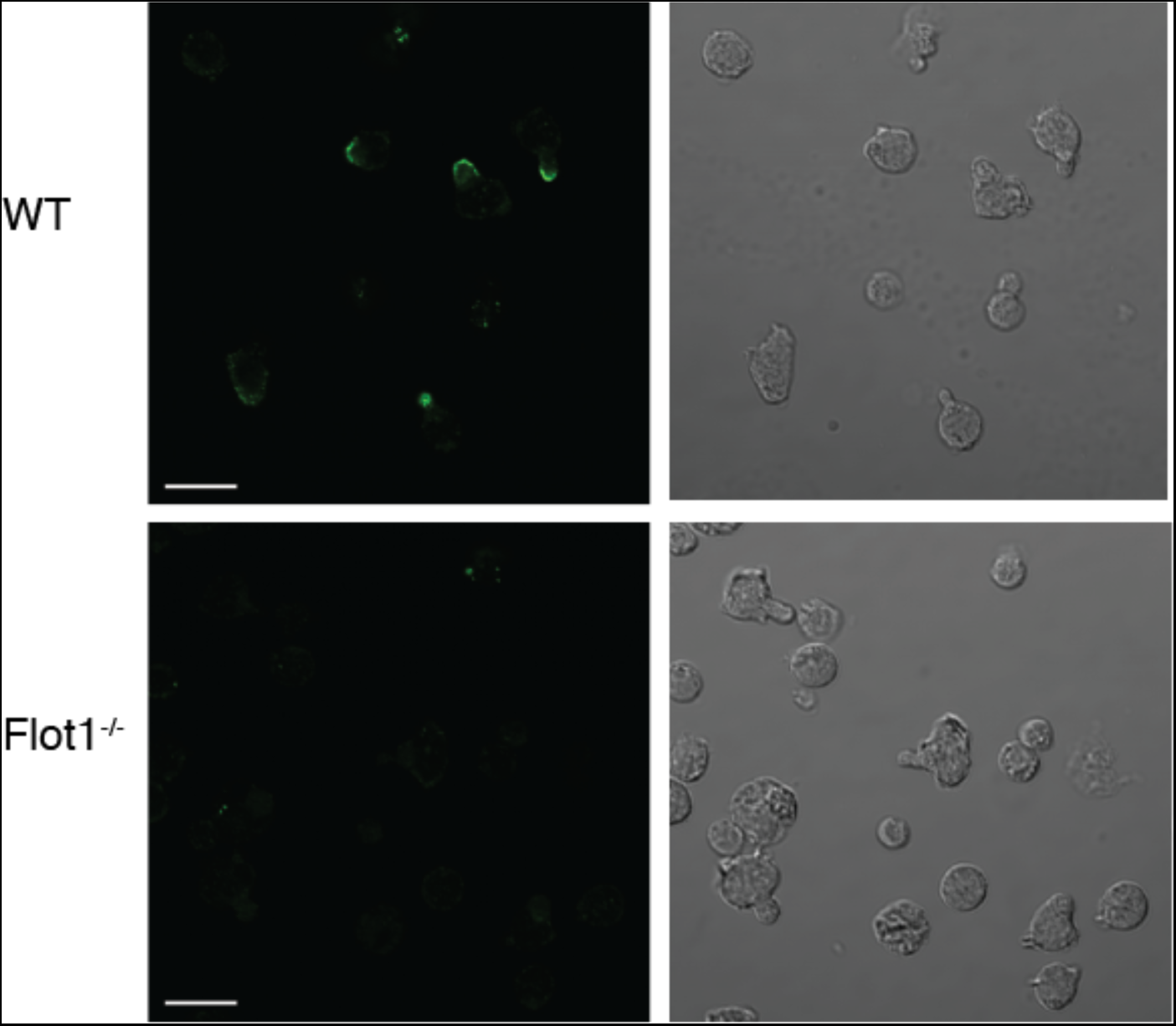
Flot1 localization at uropod of polarized T cell blasts. Phase contrast and immunofluorescent images of polarized activated WT and Flot1^-/-^ T cell blasts stimulated with CCL19 and stained for Flot1. Scale bar, 10 µm.

## Supplemental movies

**Movie S1. Naïve Flot1^-/-^ and WT OT-I T cells migrating in popliteal LN.** Scale 20 μm; Time in min:s.

**Movie S2. Memory Flot1^-/-^ and WT OT-I T cells migrating in popliteal LN.** Scale 20 μm; Time in min:s.

**Movie S3. Memory Flot1^-/-^ and WT OT-I T cells migrating in the epidermis.** Scale 50 μm; Time in h:min:s.

